# Spatially Explicit Modeling of Community Occupancy using Markov Random Field Models with Imperfect Observation: Mesocarnivores in Apostle Islands National Lakeshore

**DOI:** 10.1101/2020.08.05.238774

**Authors:** Yunyi Shen, Erik Olsen, Timothy Van Deelen

## Abstract

How species organize spatially is one of ecology’s most motivating questions. Multiple theories have been advanced and various models developed to account for the environment, interactions among species, and spatial drivers. However, relative importance comparisons of explanatory phenomena generally are neglected in these analyses. We developed a spatially explicit community occupancy model based on Markov random fields that accounts for spatial auto-correlation and interspecific interactions in occupancy while also accounting for interspecific interaction in detection. Simulations demonstrated that the model can distinguish different mechanisms of environmental sorting, such as competition and spatial-autocorrelation. We applied our model to camera trap data from a fisher (*Pekania pennanti*)-marten (*Martes americana*) and coyote (*Canis latrans*)-fox (*Vulpes vulpes*) system in Apostle Island National Lakeshore (Wisconsin, USA). Model results indicated that the observed partitioning pattern between marten and fisher distributions could be explained best by a flipped mainland-island source-sink pattern rather than by competition. For the coyote-fox system, we determined that, in addition to a mainland-island source-sink pattern, there was a positive association between fox and coyote that deserved further study. Our model could be readily applied to other landscapes (island and non-island systems), enhancing our understanding of species coexistence patterns.

## 1: Introduction

Drivers of species distributions and community structures are among the most important questions in ecology. Classical theories include niche theory (Hutchinson, 1957) and Lotka-Volterra models (Lotka, 1910; Volterra, 1928). These theories concentrated on interactions between species and their environments (hereafter: **environment sorting** where environmental filters select for species with certain characteristics and allow them to coexist) and interaction among species (hearafter:**interspecies interactions**, e.g. competition) while generally ignoring the spatial arrangement of habitat patches.

In contrast, MacArthur and Wilson (2001) emphasized the importance of random patch-level colonization and extinction probabilities in forming species richness patterns. This idea was further developed by Hubbell’s neutral theory on community assemblage (Hubbell (2001), see Volkov et al. (2003) for a review). Meta-population modelling is another example of spatially explicit theory, which emphasized the importance of dispersal (Hanski, 1983). This paradigm emphasized the importance of geographic arrangement of habitat patches (**spatial processes**, e.g. spatial auto-correlations) in determining the distributions of species. However, recent research suggested that communities reflect both species- and patch-level drivers. Leibold et al. (2004) extended meta-population models to examine community assembly while accounting for spatial and natural history processes concurrently. Yet, the relative importance of natural history processes (environmental sorting and species interactions) and spatial processes remained unclear in most communities and unaddressed in most analyses. Researchers have examined plant communities (e.g. Lasky et al. (2017)), marine systems (e.g. Shurin et al. (2009); Göthe et al. (2013); Meyer (2017)), and microbial systems using separate spatial, environmental, and interspecific interactions in both experimental and natural communities (see Logue et al. (2011) for a review). These studies suggested that a gradient from almost fully spatial-driven to almost fully environment/interaction-driven patterns in community assemblage. It is a challenge to distinguish different drivers and evaluating the relative contribution of different drivers partially due to the lack of a statistical modelling framework that accounts for these processes in a unified and comparable manner.

Markov Random Field model (MRF) defines joint distributions of sets of random variables linked by non-directed graphics (Vanmarcke, 2010; Cressie, 1992). MRF has long been used to model spatial correlations in ecology and agriculture, e.g. in spatial ecology (Hughes et al., 2011; Hepler et al., 2018), as well as temporal analysis (Zhu et al., 2005) and interspecific interactions (Harris, 2016). MRF was also widely used for modeling other types of networks (West et al., 2014; Wei and Li, 2007; Harris, 2016). Thus MRF-based models could be used as the core framework for joint modeling of environmental, interspecific interaction, and spatial drivers in a unified way so that one could examine the effect of different drivers.

Due to recent advancements in camera-trap technology, ecologists can deploy infra-red triggered cameras taking photographs or videos of passing animals that generate vast quantities of detection/non-detection data to infer presenceabsence (P/A) of moderate to large-sized animals (Burton et al., 2015) that can be analyzed using MRF models. However, imperfect detection (i.e. false absence due to imperfect sampling) remains a challenge (Kéry and Schmidt, 2008). Occupancy modelling (MacKenzie et al., 2003) addresses imperfect detection by modeling a detection process explicitly and estimating detection rates from repeated sampling.

Following a basic framework of modeling occupancy and detection hierarchically, one can build various occupancy-like models based on the idea that observations are samples taken from detection distributions conditioned on unobserved latent true patterns that follow other characteristic distributions. MRF models with imperfect observations were also explored in image reconstruction contexts (Chalmond, 1989; Ibáñez and Simó, 2003). To understand both environmental sorting and interspecific interactions, researchers proposed the use of two kinds of multispecies occupancy models. The occupancy model developed by Rota et al. (2016) used a multinomial-logistic regression which estimated different predictors for different coexistence patterns, while Kéry and Royle (2008) used a hierarchical structure to model species interactions that can be viewed as a Bayes network (Koller and Friedman, 2009). These techniques facilitated research on the assembly of animal communities in island systems and other landscapes. However, neither Kéry and Royle (2008) nor Koller and Friedman (2009) can model interactions of species and spatial processes simultaneously and explicitly in order to understand their relative importance (Cottenie, 2005; Dray et al., 2006). Too many possible patterns make multi-logistic modeling intractable as in Rota et al. (2016) and spatial autocorrelation cannot be represented using directed graphs as in Kéry and Royle (2008) while there might not exist a clear root (or dominate) species as required in Kéry and Schmidt (2008).

Our objective was to develop a model to capture spatial auto-correlation and interspecific interactions while controlling for environmental predictors. As a case study, we applied this new model to identify the drivers of species distributions for pairs of closely competing carnivore species in the Apostle Islands National Lakeshore (APIS, Wisconsin, USA). Since the case study is an island system, we also tested whether the distribution patterns followed a mainland-island pattern or a stepping stone pattern.

Notably, island systems are a special case where spatial processes might play an outsize role, despite this we intend for our model to be applicable to the island and non-island systems. We anticipate this method could be applied in any situation where researchers would like their models to explicitly account for predictors, spatial auto-correlation, and inter-specific interactions in a unified framework. For instance, a camera trap survey where the grid is small enough to impose spatial auto-correlation.

## 2: Methods

### 2.1. Study Area

Apostle Island National Lake Shore (APIS) is located on the southwest shore of Lake Superior (USA) and lies in the transition zone between temperate and boreal forest regions. APIS is distinct from tropical islands (where much research on community assembly has occurred) because of severe winters (84.6 days for maximum and 185.4 days for minimum temperature below 0°C while 29.4 days minimum temperature below −17.8°C, Arguez et al. (2010)) and relatively low primary productivity (average net primary productivity of 2015 is 8.23 × 10^3^kgC/m^2^/year, Running and Zhao (2018)). Ten species of native carnivores were detected during 2014-2017 (Allen et al., 2017). How these species coexist and how richness differs among islands is a fundamental question for understanding community dynamics in temperate island systems.

### 2.2. Camera trapping surveys

During 2014-2017, APIS staff and collaborators conducted camera-trap surveys to determine distributions and relative abundances of mammals within the National Lakeshore (Allen et al., 2017). Twentyone of 22 islands that make up the archipelago were surveyed using a 1 km^2^ lattice (grid) sampling frame. Within each grid cell, there was one camera trap. Scent lures were placed at half of the camera stations at deployment and rotated to the remaining stations during mid-deployment camera checks. Since all islands could not be monitored at the same time, though substantial overlap did occur, we assume the underlying distribution of species did not change during the 3-year survey period. We divided surveys into 60-day blocks to create repeat observations. The model allowed us to incorporate predictors to account for differences in detection during periods however we were not able to obtain useful predictors for all periods and sites because of incomplete environmental data.

### 2.3. Hypothesis

We focus on two pairs of plausibly competing species: fisher (*Pekania pennanti*)-marten (*Martes americana*) (fisher-marten system) and coyote (*Canis latrans*)-red fox (*Vulpes vulpes*) (coyote-fox system). In APIS, 30% of sites had fisher detections also had marten detections, and 15% of sites that had marten detections also had fisher detections (2014-2017), in contrast, 64% of sites that had red fox detections also had coyote detections, and 28% of sites that had coyote detections also had red fox detections (Fig.1). In niche-based theories, a partition pattern like fisher and marten could be understood as spatial niche partitioning due to competition, while coexistence could be achieved by partitioning other niche dimensions such as time. In these theories, competition was a factor promoting the partition pattern. In dispersal-related theories, species were more likely to exist close to the source of dispersing individuals which is usually a stable mainland population. If species were independent, they would coexist on islands closer to the mainland with a higher probability subject to dispersal capability. Since we observe a spatially partitioned pattern for fisher-marten, it may be explained by competition or different source-sink dynamics (spatial processes). Importantly, we observed that marten tended to occur on more distant islands that might indicate a “flipped” mainland-island pattern opposite the prediction of the mainland as the source. For coyote and fox, coexistence at islands closer to the mainland could be explained by dispersal from the mainland. Yet, after accounting for such potential spatial processes, we are interested also in determining if these two species associate or compete (dominant competitor excludes sub-dominant). The environment across the islands is rather homogeneous thus we were unable to obtain useful environmental predictors, hence, the only “environmental” predictor considered here was the distance to the mainland (negative exponential transformed) which should be considered as part of a spatial process.

**Fig. 1.**
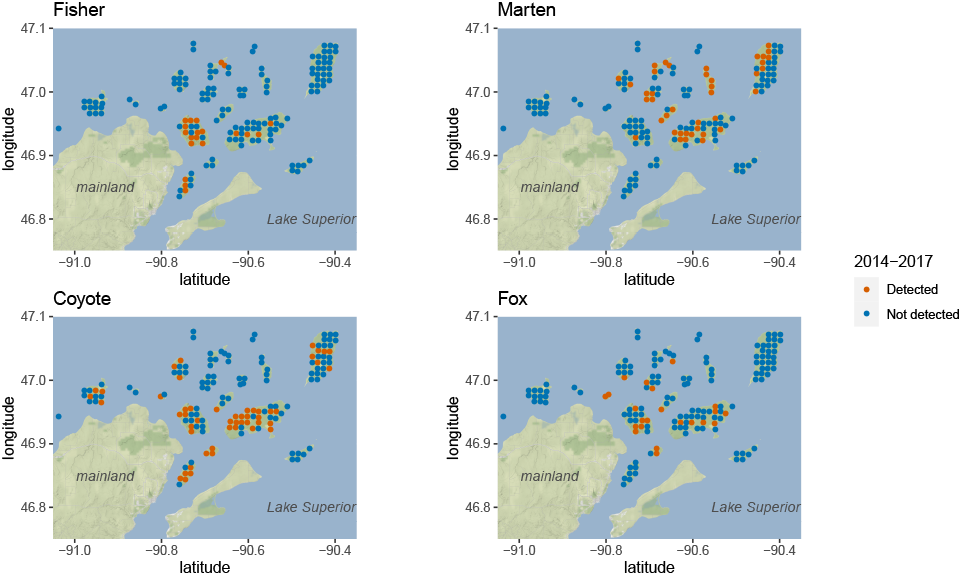
Detection of 4 target species on the islands, blue: detected, red: not detected, dots represent camera locations on the APIS

**Fig. 2.**
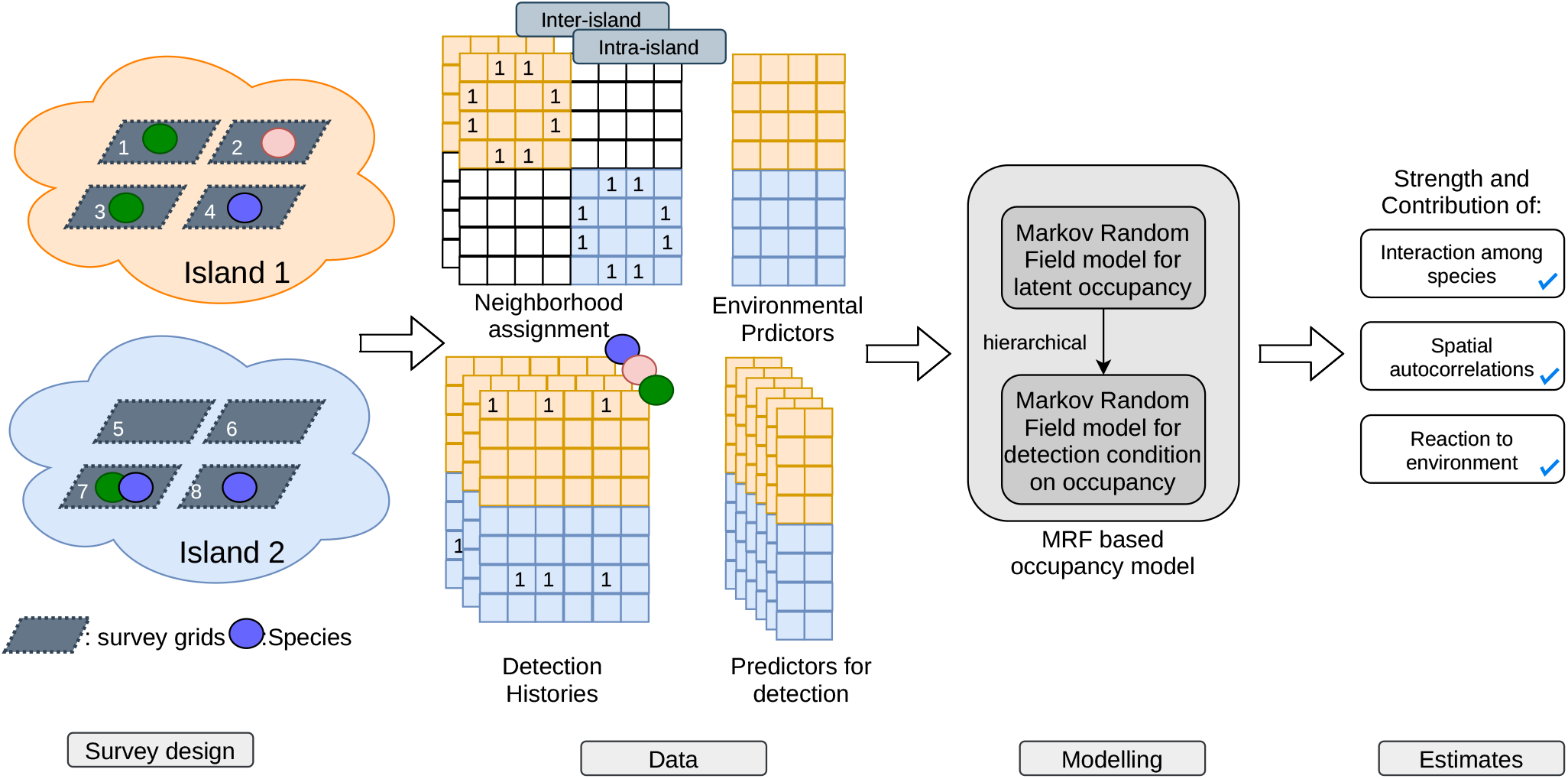
Workflow of the modelling. The flow chart represents the data requirement and structure of the model. Our case study was done in an island setting but note that any grid system can be viewed as a single island system while a multi-season system can be viewed as having multiple “temporal islands”.

For the fisher-marten system, we pose two working hypotheses for the observed pattern:

1. Distribution of both species among the islands reflects similar mainland-distance dependencies (spatial effects). While separation at the site level within the island is due to competition (interaction effects)
2. Distribution of both species among the islands reflects differing mainland-distance dependencies but shows minor within island competition at the site level.

For the coyote-fox system, we pose two working hypotheses for the observed pattern:

1. Distribution of both species in the islands reflects spatial drivers. Coexistence is facilitated by separation in time.
2. Trophic position and life-history drives distribution (foxes avoid coyotes at the site level), while spatial effects at the island level are minor.

Besides comparing strengths of different drivers, we also sought to test whether the spatial auto-correlation followed a mainland-island model where there are no spatial auto-correlations between sites on different islands while the distance to mainland played an important role or a steppingstone model where there are spatial auto-correlations between sites on different islands that are close and mainland mainly affects the closest islands.

### 2.4. Spatial Explicit Community Occupancy Analysis using Binary Markov Random Field Model

The analysis was implemented in R (R Core Team, 2019) while the core algorithm to fit the proposed model was partially implemented in C++ with R interfaces (using Rcpp and RcppArmadillo Eddelbuettel and Sanderson (2014); Eddelbuettel and François (2011)). Implementations can be found on the author’s GitHub repository YunyiShen/IsingOccu-core.

We used an Ising model (Ising, 1925), a kind of Markov Random Field (MRF) model whose responses are all binary, to model the distribution of competing species in a spatially explicit manner. Coding of latent true occupancy status followed the convention in network science, i.e. +1 for presence and −1 for absence. This symmetric coding was more conventional in physics but less so in ecology. Models with a centering term mostly used in ecology enabled modeling of the “large scale” response due to environmental predictors (centered auto-logistic models Hughes et al. (2011)) which tried to detect auto-correlation in the residuals of a large scale response due to environmental drivers. However theoretical studies with this model by Wolters (2017) suggested better performance of a symmetrically coded model compare to a centered model when associations (e.g. competition, spatial autocorrelations) were expected. Additionally, centered models predicted non-linear relationships between the strength of association and log odds of two species coexisting or co-absenting while symmetrical coding did not have this problem. Further, symmetrical coding avoided the cross-product between different terms (i.e. environment and species interactions) in the negative potential function (log probability mass function (pmf) up to a constant difference) of this model (Koller and Friedman, 2009) which helped us evaluate relative contributions of different mechanisms. Most importantly, parameters of a symmetrical-coding model had better conditional interpretation, e.g. regression coefficient (*β*s), interaction terms (*γ*s), and auto-correlation strength parameters (*η*s) which are the conditional log odds of presence given all other sites and species while other multispecies/spatial-explicit frameworks (e.g. Rota et al. (2016); Hughes et al. (2011)) do not have this clear interpretation. This property was important in understanding the strength of different mechanisms since interactions should be conditioning on environments, while the reaction to the environment should also be conditioning on interactions e.g. Blanchet et al. (2020), argument 2 for instance.

To make comparisons between environmental sorting, spatial processes, and interspecific interactions as drivers of species’ spatial distributions, three components were considered simultaneously but separately in the graph associated with the joint distribution: 1) a linear predictor calculated from environmental covariates, 2) a nearest neighborhood spatial autocorrelation at camera-site level (site level hereafter,Hepler et al. (2018)) within and among islands, and 3) local species associations at the site level (We assume that partial associations reflect interactions, similar to Harris (2016)). We denote the design matrix for environmental covariates as **X** (assumed fully known and of full column rank) and responses of certain species *k* (*k* = 1, 2.., *w*) to environment **X** as *β*_*k*_. Further in this case study, due to the different nature of site linkages within and across islands, inter-island and intra-island correlations were modeled separately. We denote the strength parameters of these two correlations as *η*^*ex*^ and *η*^*in*^, and known adjacency matrix **D**^*ex*^, **D**^*in*^ (eqn.1). Mainland-island with linkage matrix *D*^*ml*^ shares the same strength of inter-island spatial autocorrelation in this study. We denote the presence and absence vector of species *k* on the landscape as **Z**_*k*_. We denote the transpose of **Z**_*k*_ as 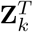 Thus, the joint distribution of all species at all sites has the form:

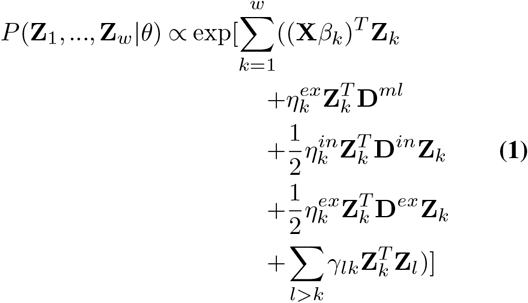

For the detailed meaning of parameters see Table.1. We use *θ* as an abbreviation of all parameters in a conditional probability. Note that the first term accounts for an environment response (mainland-island effect), the second account for mainland-island processes (as a special environment predictor, mainland-island spatial effect), the third term accounts for intra-island spatial auto-correlations (spatial effect), the fourth term accounts for inter-island spatial auto-correlations (and can be other types of auto-correlations) and the last term accounts for all inter-specific interactions between species *k* and species *l* while *γ*_*lk*_ is the strength of association between the two species. In the mainland-island setting, we assumed that there was no inter-island spatial autocorrelation so **D**^*ex*^ has all 0 as its entries.

**Table 1.**
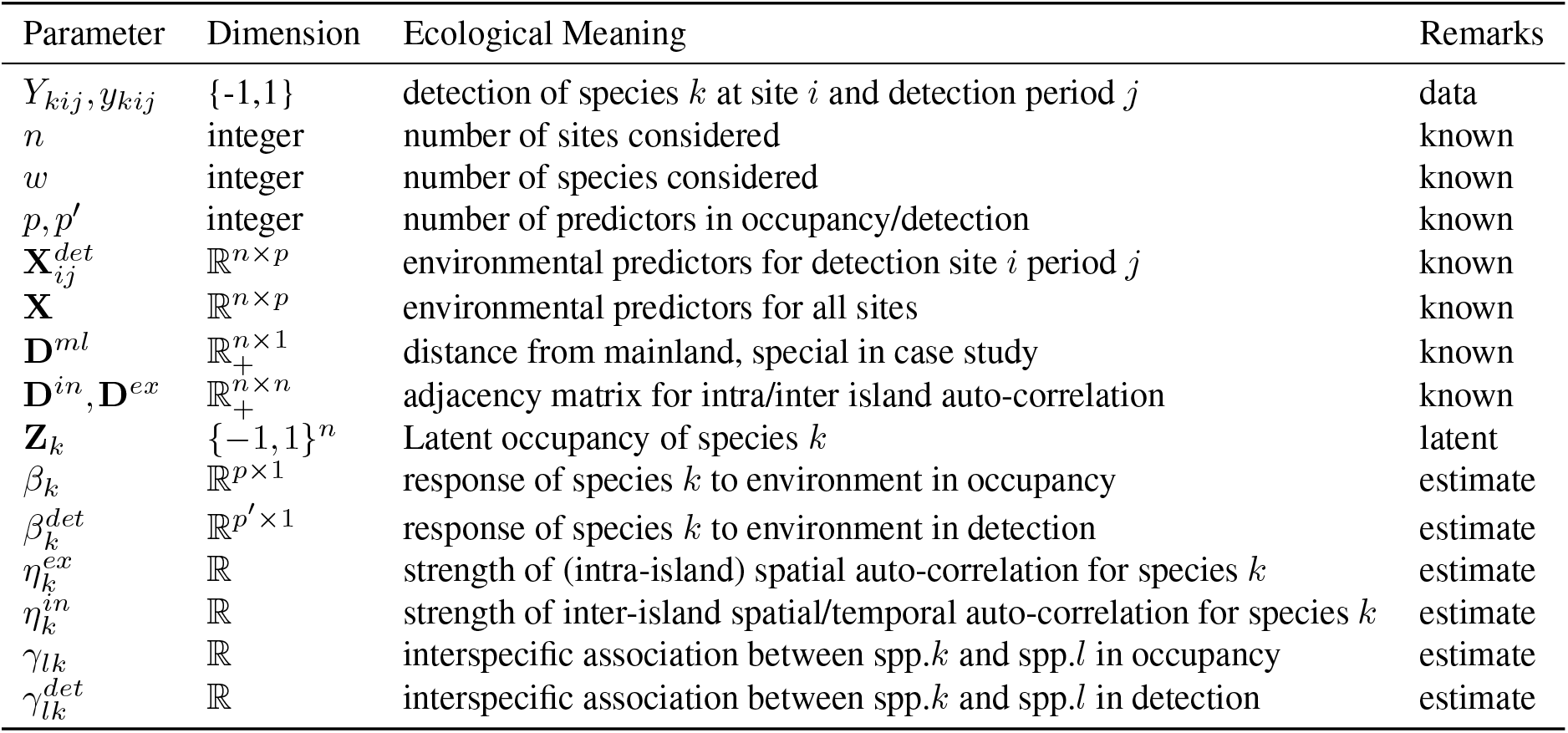
Model Parameters.

### 2.5. Accounting for Imperfect Detection and Short-term Interactions

Following the logic of occupancy-like modeling (MacKenzie et al., 2003), we modeled observed detection/non-detection as repeated samples from a detection process. Associations in short-term detection can also be informative about species interaction. We further assumed that the interspecific interactions are local (i.e. no spatial auto-correlations considered in the detection process). We used another binary MRF (Ising model) conditioned on the occupancy status of a species to model the detection process. In total, there were two binary MRF models: 1) latent occupancy, and 2) detection conditioned on occupancy. Only species occupying a certain site were included in the detection MRF and species not occupying had a probability of non-detection of 1. Formally, we denote *y*_*kij*_ as species *k*’s detection status at site *i* during period *j*. The detection model also included environmental predictors of each site for a detection period with the design matrix denoted by 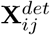 (assumed fully known and of full column rank) with the regression coefficient of species *k* in detection part 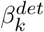. The likelihood function at site *i* and detection repeat *j* is given by eqn.2.

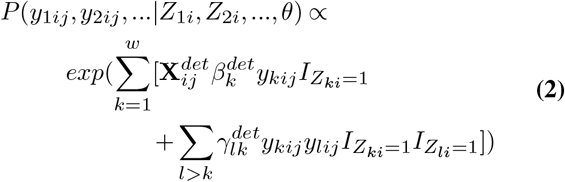

We used *θ* as an abbreviation of all other parameters in the conditional probability. *I*_*{}*_ is the indicator function and 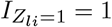 only if *Z*_*li*_ = 1 and 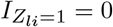 otherwise. The indicator function “knocks out” the species from detection interaction if it was not occupying that site. The reasoning behind this “knocking out” was that we assumed that non-detection was caused by the absence of a species and, thus, should be understood as a do-calculus (Pearl, 1995) rather than conditioning. Unlike the occupancy part, this conditional likelihood function is tractable for a reasonable number of species (e.g. *<* 10) because of the relatively small size of the underlying graph. The joint likelihood function of the whole detection history, conditioned on occupancy, was the product of each site and detection period. The joint (unnormalized) likelihood function of observed detection data then could be calculated by multiplying eqn.1 and eqn.2. The latent **Z**’s could be estimated similarly with unknown parameters.

Priors were set to be vague normal distributions due to the relatively small number of repeats and lack of environmental variation in our APIS case study. We put a normal prior with variance 0.1 on intercept of detection (0.95 HDR for detection rate: [0.22,0.78]) as part of our assumptions. Again this was not necessary for the model per se (as seen in simulation), but part of the case study. Sensitivity analysis on this part was also conducted. Posterior distributions were simulated through a Markov chain Monte Carlo (MCMC) algorithm (Hastings, 1970) that is often used in complex ecologically modelling (Rempel, 2011; Boulange et al., 2017). To overcome the double-intractable nature of the posterior (Murray et al., 2012; Møller et al., 2006), we followed the single parameter change method proposed by (Murray et al., 2012). We calculated a p-value level that credible interval (CI) spans 0. A full description of the algorithm used is in Appendix.S2. Diagnostic evaluation of MCMC results were done using R package `coda` (Plummer et al., 2006).

### 2.6. Selection Between Stepping-Stone and Mainland-Island Model

We compared two general models for spatial auto-correlation between islands in this study.

1. A *stepping-stone model* assumed that sites at the edge of an island can be neighbors to sites on another island in a MRF sense. We assign this linkage using Delaunay triangulation (Okabe et al. (2009), Fig.S1). The strength of correlation was assumed to decay exponentially through the normalized distance (Shurin et al., 2009). Sites on the closest islands have linkage to the mainland and the log odds of having species occupying such sites decay exponentially through the normalized distance to the mainland (Shurin et al., 2009).
2. A *mainland-island model* assumed that sites on different islands were conditionally independent given their distance to the mainland, the log odds of having species occupying a site decayed exponentially through the normalized distance to the mainland (Shurin et al., 2009).

Bayes Factor (BF), a Bayesian generalization of a Likelihood ratio test, was used for model selection (Gelman et al., 2013). We assumed the two models were equally plausible and thus the Bayes factor can be understood as the ratio between the posterior probabilities of the two models. We can calculate the posterior predictive distribution of data following Raftery et al. (2006). One obstacle to using BF in this analysis is the intractable likelihood function preventing us from directly calculating the predictive probability of each model by calculating the likelihood function during the posterior sampling. However, we could follow Descombes et al. (1999) in calculating the likelihood function by sampling augmentation variables (Appendix. S2). Since the ratio is also estimated, robustness diagnostics followed Descombes et al. (1999).

### 2.7. Evaluating Contribution of different Drivers

Different processes can drive the degree of spatial partitioning among species, e.g. environmental sorting can promote coexistence when species need similar resources (Grinnell, 1917; Saporetti-Junior et al., 2012) while competition promotes partitioning (Tilman, 1985; Gastauer and Meira-Neto, 2014). Compared with the strength and direction of each process, a different question was: which driver makes the *observed* pattern likely? For instance, is it possible that while species A and B compete, we observe that they still coexist because of environmental sorting? In this case, we may argue that environmental sorting had a larger contribution to the observed (coexistence) pattern.

We used negative potential functions (a.k.a. Hamiltonian functions in our specific setting from Cipra (1987) eqn.1.1, Osogami (2017); Pfeuty (1970)) as statistics to evaluate the contributions of each driver on the observed pattern since drivers were modeled explicitly in the proposed model. In our setting, the Hamiltonian function was the log of the right-hand side of eqn.1. These types of statistics were used to quantify “fitness landscapes” of amino acid interactions in protein systems (Levy et al., 2017; Shekhar et al., 2013; Ferguson et al., 2013; Morcos et al., 2014) with many other applications in quantifying the stability of interactions (Ezaki et al., 2017; Becker and Karplus, 1997; Cipra, 1987).

Statistically, a negative potential function can be viewed as a fitness score defined by log probability mass function (pmf) up to a constant difference. Patterns with higher scores had higher probabilities of occurrence (Levy et al., 2017). In model specifications, we used several distinct terms to account for different ecological processes and we used the score of each of the terms to evaluate the relative contribution of each process to the observed pattern. For instance, 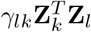 is the score for interspecific interaction between species *l* and *k* where *γ*_*lk*_ is the strength of the interaction and **Z**_*k*_ is the vector of the occupancy of species *k*. The score is an “accumulated” strength through occupancy. Note the score assesses the probability mass of the observed pattern, i.e. a high positive contribution to the negative potential function means the corresponding term made the probability mass on the observed pattern high. The question to be answered here is whether *a certain process (e*.*g. spatial) had a large contribution to the observed pattern* as we argued above. The strength of a process is different from its contribution to the observed pattern, a different aspect of the hypothesis. Posterior distributions for different terms of negative potential functions were calculated using posterior samples of latent occupancy status and model parameters.

### 2.8. Simulations

We conducted simulations using the same spatial arrangement of APIS’ camera trapping grids, as well as regular 10 ×10 (i.e. 10 rows and 10 columns of sites), 15 ×15, 20 ×20, and 25 ×25 grids. On APIS’ camera grids, we tested three mechanisms: 1) Competition, 2) Non-interaction, and 3) Sorting. In competition simulations, two species had the same reaction to the environment (or distance to the mainland for APIS) and a negative association. In no-interaction simulations, two species had opposing reactions to the environment or distance to the mainland (represent niche difference/flipped source-sink) and no association. In sorting simulation, two species had the same but relatively weak reaction to the environment or mainland distance and a positive association. There were spatial auto-correlations on all mechanisms. On regular grids, we tested two species on a random draw landscape with one environmental predictor (note that we did not randomize this environment) while on APIS, we used distance to the mainland as the predictor since there was a lack of environmental variation. For detailed simulation settings see Table.S1 and Table. S2. In simulations, all parameters had vague priors.

### 2.9. Posterior Predictive Checks

To check for systematic discrepancies between data and model predictions, we performed posterior predictive checks (Gelman et al., 1996; Lynch and Western, 2004; Gelman et al., 2013) using 6 metrics for the APIS case study. They were: 1) frequency of detection for fisher and marten and 2) frequency of detection for coyote and fox, this represented how often we see the species; 3) proportion of sites that had at least one detection for fisher and marten or 4) for coyote and fox, this represented the naive occupancy rate of 4 species; 5) correlations between naive occupancy in each system (fisher-marten or coyote-fox), in which naive occupancy was 1 if species were ever seen at that site and -1 otherwise; 6) count of sites that had both species from either of our species-systems detected. In total 2,000 posterior predictive detection histories with the same time frame as the original APIS data were sampled and posterior predictive p-values for each statistic were calculated. Small p-values indicated variations the model failed to capture.

## 3: Results

### 3.1. Simulations

We evaluated differences between posterior median estimates and true parameter values in 100 simulated datasets (Fig.3). In all cases, the middle 50% quantile contained 0, suggesting that we generally could recover parameter values using regression models. However, we were conservative regarding spatial auto-correlations (posterior medians were less than true parameter values in some cases) due to relatively a small number of grids, as also shown in Hughes et al. (2011). When there are no repeats (i.e. we have only one occupancy sample), we could reduce the uncertainty on spatial auto-correlation strength by having larger grids.

**Fig. 3.**
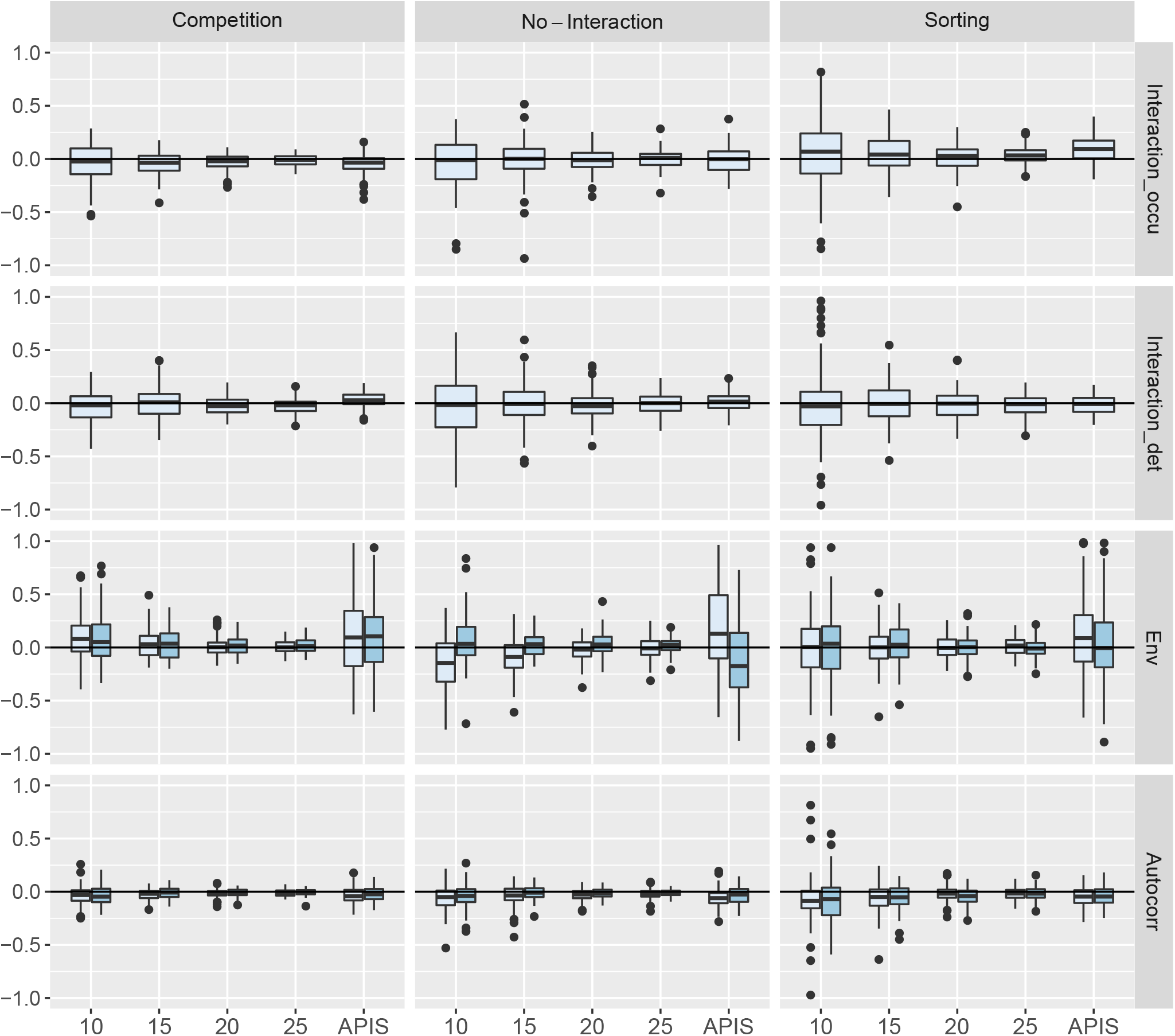
Differences between posterior medians and true parameter values of 100 simulated data sets. First column: *Competition*, species had same environment dependency and negative interspecific interaction, Second column: *No interaction*, species had different environment dependency and no interaction, Third column: *Sorting*, species had same environment dependency and positive interspecific interaction. Rows correspond to parameters estimated in the model. First row: Interaction in occupancy, Second row: Interaction in detection, Third row: Reaction to the environment, Fourth row: Spatial autocorrelation. The x-axis was the size of lattice or APIS (155 grids). Shading of boxes indicates each of the two “species” simulated, note that interaction parameters are between species thus no shading difference. Boxes represent quartiles.

**Fig. 4.**
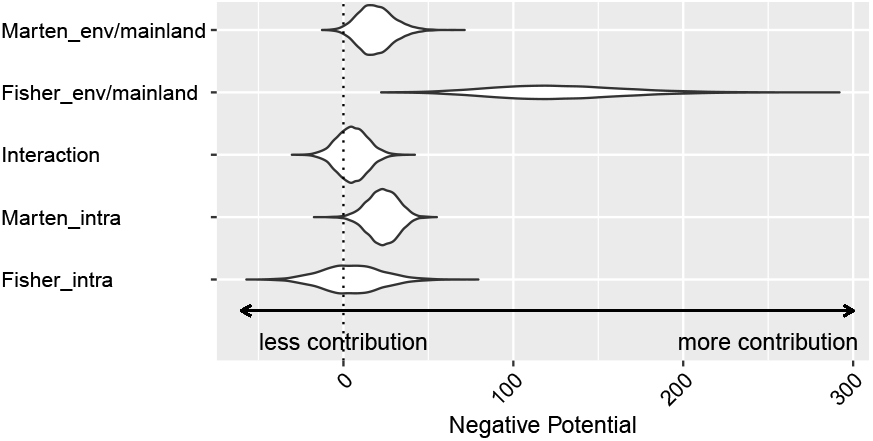
Posterior distribution of different terms in negative potential function for the Fisher-Marten system. Values represent the contribution of a certain term in the negative potential function, higher value represent a higher contribution to the observed occupancy pattern, note that we combined mainland-island and intercept which represented the overall environment.

### 3.2. Fisher-Marten System

A total of 3 ×10^6^ samples were drawn after 5 ×10^4^ burn-in and thinned by 300. Diagnostics of MCMC showed sufficient mixing of the chains (Fig.S2), suggesting that the MCMC algorithm can approximate the posterior distribution of parameters. Log_10_ Bayes factors for mainland-island models and stepping stone models were estimated to be 4.26, (i.e. the posterior probability of mainland-island model was 10^4^ higher than the posterior probability of stepping stone model) hence data decisively supports mainland-island rather than the stepping stone model following the recommended cutoff of 2 (Kass and Raftery, 1995). This result suggested that the spatial pattern of the system can be explained better by a mainland-island model rather than a stepping stone model. Further analysis was based on the Mainland-Island model.

Table.2 shows posterior estimates of model parameters of interest for the mainland–island model of the fisher-marten system. We detected a significant positive distance dependency (positive means a higher chance of occupying a closer island here because the distance is negative exponentially transformed) in fisher (*η*^*ex*^ = 2.479, *CI* = [0.976, 4.744]) and a negative distance dependency in marten occupancy (*η*^*ex*^ = *−*0.789, *CI* = [−1.907, *−*0.0558]).

**Table 2.**
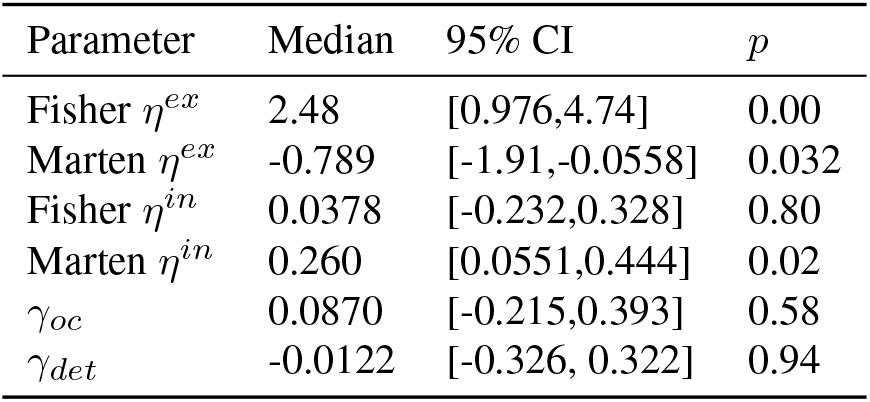
Posterior estimation of model parameters in Fisher-Marten mainland-island system. *η*^*ex*^ represented the distance dependency, *η*^*ex*^ *>* 0 meant decay through distance, *η*^*in*^ represented the intra-island spatial auto-correlation, *γ*^*oc*^ represented the association between species in occupancy and *γ*^*det*^ represented the association between species in detection, p is the level that CI cross 0

| Parameter | Median | 95% CI | p |
| --- | --- | --- | --- |
| Fisher $\eta^{ex}$ | 2.48 | [0.976, 4.74] | 0.00 |
| Marten $\eta^{ex}$ | -0.789 | [-1.91, -0.0558] | 0.032 |
| Fisher $\eta^{in}$ | 0.0378 | [-0.232, 0.328] | 0.80 |
| Marten $\eta^{in}$ | 0.260 | [0.0551, 0.444] | 0.02 |
| $\gamma_{oc}$ | 0.0870 | [-0.215, 0.393] | 0.58 |
| $\gamma_{det}$ | -0.0122 | [-0.326, 0.322] | 0.94 |

However, we did not detect a significant association between these species in either occupancy or detection (*CI* = [−0.215, 0.393], [−0.326, 0.322]). Marten showed an intra-island spatial auto-correlation 0.260 *CI* = [0.0551, 0.444]). This result supported the second hypothesis that fisher and marten had a “flipped” mainland-island pattern whereby fisher had a higher chance of occupying closer islands while martens had a higher probability of occupying more remote islands independently of possible competition.

As discussed in our methods, the strength of a certain factor is a different question compared to the contribution of the factor. Posterior distributions of negative potential functions in the fisher-marten system suggested that association is weak and had no significant contribution to the observed distribution pattern of fisher and marten. The fisher-marten system seemed to be dispersal/environment driven for fisher, while both dispersal/environment and intra-island spatial auto-correlation had similar levels of contribution for marten. Hence, competition is weak and had a minor contribution to the observed marten pattern which indicated a flipped mainland-island hypothesis where the mainland functions as a sink rather than a source.

Posterior predictive checking showed all 6 p-values were greater than 0.05 indicating no systematic discrepancies between data and model predictions which suggested fitness of the fitted model (Fig.5).

**Fig. 5.**
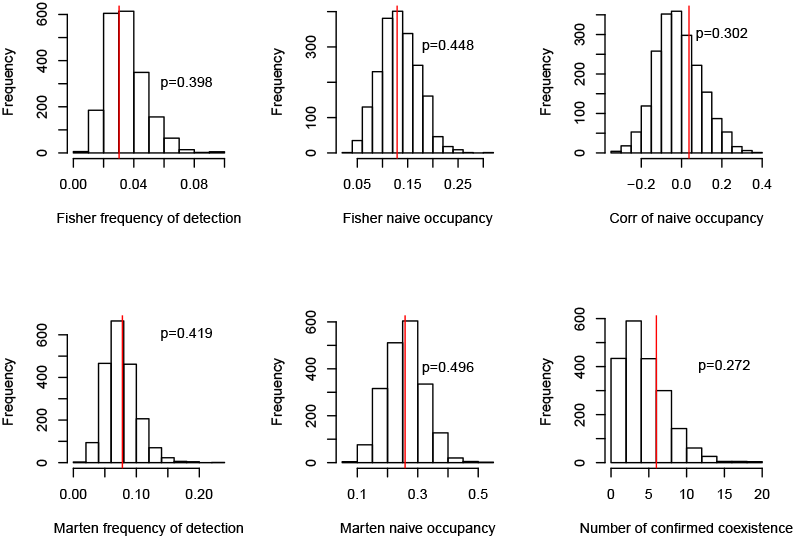
Posterior predictive checking for the Fisher-Marten system and assessing model fitness. All p-values are large suggested that there were no conflict between data (red line) and predictions of our model according to the 6 statistics used thus a good fitness of the model.

### 3.3. Coyote-Fox System

Log_10_ Bayes factors (log posterior odds of two models) for mainland-island and stepping stone models were estimated to be 18.3 (i.e. the posterior probability of mainland-island model was 10^18^ higher than the posterior probability of stepping stone model), hence data decisively supported the mainland-island rather than the stepping stone model following the recommended cutoff of 2 (Kass and Raftery, 1995). This result suggested that the spatial pattern of the system can be explained better by a mainland-island model rather than a stepping stone model.

Further analysis was based on the Mainland-Island model. Posterior estimates of model parameters (Table.3) indicated a significant positive distance dependency in fox but not coyote (Coyote:*η*^*ex*^ = 0.552, *CI* = [−0.378, 1.69] Fox: *η*^*ex*^ = 2.41, *CI* = [0.428, 6.30]) (Table.3). Meanwhile, posterior association in occupancy was estimated as positive but only weakly significant (*γ*_*oc*_ = 0.234, *CI* = [−0.041, 0.53]. The posterior probability of this parameter being positive was *p*(*γ*_*oc*_ *>* 0|*data*) = 0.95), suggesting that despite dispersal/environment drivers, there may be evidence of a positive association between two species at grid level that needs further evaluation. These findings supported the hypothesis that the coyote-fox system might not be fully explained by spatial factors and require further evaluation. Notably, we detected a significant positive association in detections (*γ*_*det*_ = 0.427, *CI* = [0.211, 0.646]) which might suggest a behavioral association.

**Table 3.** Posterior estimates of model parameters in Coyote-Fox mainland-island system. *η*^*ex*^ represented the distance dependency, *η*^*ex*^ *>* 0 meant decay through distance, *η*^*in*^ represented the intra-island spatial auto-correlation, *γ*^*oc*^ represented the association between species in occupancy and *γ*^*det*^ represented the association between species in detection, p is the level that CI cross 0

| Parameter | Median | 95% CI | p |
| --- | --- | --- | --- |
| Coyote $\eta^{ex}$ | 0.552 | [-0.378 1.69] | 0.24 |
| Fox $\eta^{ex}$ | 2.41 | [0.428 6.30] | 0.02 |
| Coyote $\eta^{in}$ | 0.196 | [-0.0186 0.426] | 0.08 |
| Fox $\eta^{in}$ | -0.0696 | [-0.345 0.207] | 0.60 |
| $\gamma_{oc}$ | 0.234 | [-0.0411 0.528] | 0.1 |
| $\gamma_{det}$ | 0.427 | [0.211 0.646] | 0.00 |

Similar to the fisher-marten system, dispersal/environment also had the largest contribution to the observed pattern in the coyote-fox system (Fig.6). However, in contrast with fisher-marten, both spatial auto-correlation and interspecific interaction seemed to have some importance (Fig.6) as also suggested by the hypothesis that associations exist that are additive to spatial drivers.

**Fig. 6.**
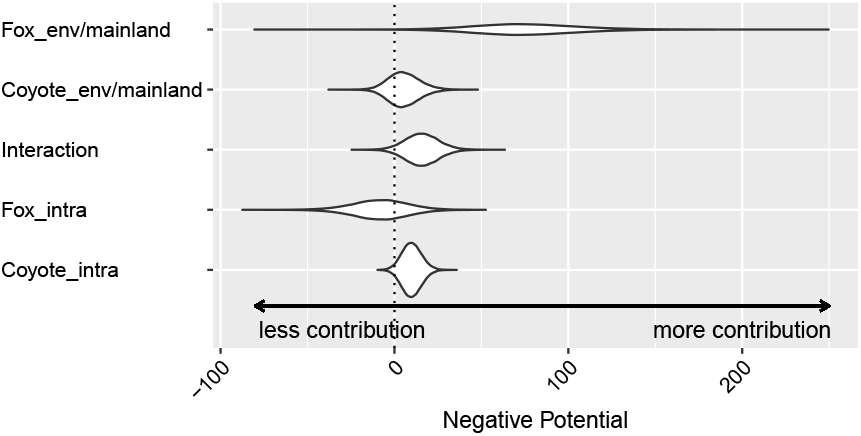
Posterior distribution of different terms in negative potential function for the Coyote-Fox system. Values represent the contribution of certain terms in the negative potential function, higher value represent a higher contribution to the observed occupancy pattern, note that we combined mainland-island and intercept which represented the overall environment.

Posterior predictive checking showed all 6 p-values were greater than 0.05 thus no systematic discrepancies between data and model predictions which suggested the fitness of the fitted model (Fig.7).

**Fig. 7.**
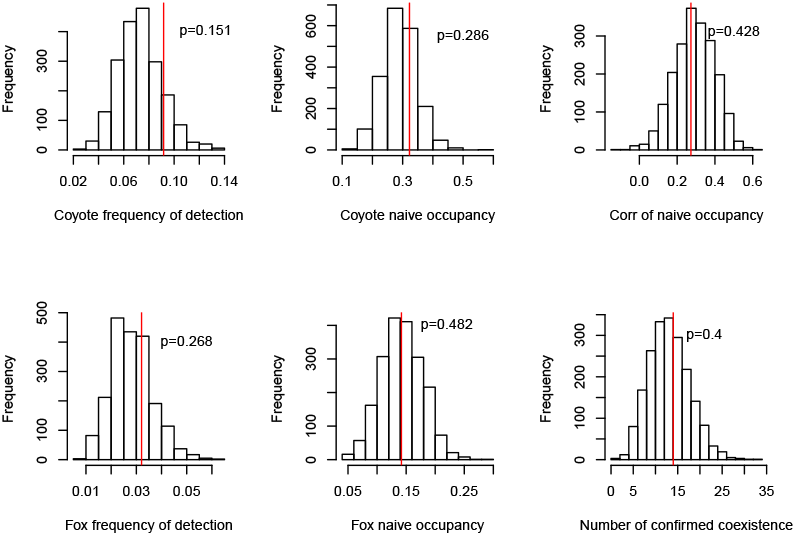
Posterior predictive checking for the Coyote-Fox system assessing model fitness. All p-values are large suggested that there were no conflict between data (red line) and predictions of our model according to the 6 statistics used thus a good fitness of the model.

## 4: Discussion

We developed and tested a MRF-based, multispecies, spatially explicit, occupancy model which allowed the evaluation of relative contributions of spatial auto-correlations and life history mechanisms in influencing the coexistence pattern of species. Our model enables ecologists who conduct research on community structure to consider spatial and life history drivers jointly and explicitly. Though the model assumed patterns remain constant through time (single season), extending it to a multiseason analysis is straightforward. For instance, within our proposed model framework, one could define “sites” as spatial sites at a certain time, then by applying an adequate adjacency matrix (e.g. **D**^*ex*^) so that only the individual spatial sites at adjacent season were connected. By doing this one can incorporate temporal auto-correlations similar to spatial auto-correlations given we can have two or more sources of auto-correlation in the proposed framework. Similar comparisons of the strength and contribution can carry through in a multiseason analysis. Another way of viewing this is by viewing season as temporal islands and we can have a stepping stone model on time. To incorporate extinction and colonization, one could use the dynamic Ising model as the core (Nguyen et al., 2017) in the place of the static Ising model used here. Compared with Bayesian network-based multispecies frameworks proposed by Kéry and Royle (2008), our method did not ask for a species to be the root of the network and allowed cycles in the network. Moreover, analysts can condition occupancy on another species to accommodate a single dominant competitor. Compared with Rota et al. (2016), our method had a better interpretation especially when a species network was large; since, in our method, the interaction between species was modeled explicitly by auto-regression terms. Neither Kéry and Royle (2008) nor Rota et al. (2016) were spatially explicit. Partly because the graph represented the spatial correlation, it had no natural direction and could not be represented by a directed graph like Kéry and Royle (2008) did for species. Also when the number of potential outcomes was too large one cannot assign unique linear predictors for each pattern as in Rota et al. (2016). Markov random field modeling was used in quantifying interspecific interactions in Harris (2016) and can help identify interactions between species when controlling for the environment and other confounding interactions (e.g. apparent competition where A, B both interact with C while no interaction occurs between A and B) (Blanchet et al., 2020). Again, though our case study is an island system, the model is not specific to island systems and the case study did not demonstrate the full potential of the model (and software) we built (e.g. our method also allows for the inclusion of predictors in detection). Since the explicit terms of environmental predictors, spatial autocorrelations of different types, and interspecific interactions can all be realized with this method, one could apply this model to any system where these drivers should be considered and compared in a unified way. For instance, habitat patches, or camera trap survey grids.

Our results on two pairs of plausibly competing species as components of the meso-carnivore community also showed that community structure reflected drivers associated with two broad theoretical paradigms. First, we detected positive intra-island spatial-autocorrelation for 2 species out of 4. This spatial autocorrelation term makes sites no longer exchangeable even when the distance to the mainland was controlled. Spatial autocorrelation also had different strengths for the four species considered, which means species were not exchangeable even when considering spatial processes. Coyotes and foxes had different strengths of dependence on mainland distance, likely due to different dispersal ability, i.e. coyotes (which are larger) likely can disperse farther than foxes and, thus have weaker distance dependency.

We detected an unexpected opposite direction of mainland distance dependency on the occupancy of marten. The fisher-marten system provides a possible example where the mainland serves as a sink rather than a source for a species from a metapopulation point of view. This spatial pattern was recently identified independently for martens using genetic analysis (Smith et al. (2020), in press). A more general meta-community framework should be used when considering island or island-like systems. If two species follow similar dispersal patterns but need to partition spatially we should expect closer islands to be more likely occupied by one of the species than the further islands and partitioning should happen on islands with similar, nearby distances to the mainland (i.e. perpendicular to the mainland-island dispersion direction). However, we observed similar levels of *co-absence* on islands regardless of their distances to the mainland, i.e. martens were not occupying nearby islands regardless of fisher presence. Meanwhile, the partitioning of fisher and marten happened in parallel (distance dependency) rather than independently of distance (i.e., competition) to the mainland-island dispersal direction which was different from what we would expect based on hypothesis 1 (i.e., distance dependency with competition). We did not observe fisher and marten occupying sites closer to the mainland or partitioning at the site level. The fisher-marten system appeared to better conform to hypothesis 2 because fisher dispersal direction appeared to be from the mainland to the islands, while marten dispersal appeared to be from the islands to the mainland. However, these results were solely from distribution data and additional evidence from genetics (e.g., Smith et al. (2020), in press), movement data, or behavior, etc. would be needed to further support this argument.

For the coyote-fox system, we observed that spatial autocorrelation had a strong influence on coyote distribution. This may be due to the relatively small size of the islands compared to coyote home ranges. Typical coyote home ranges were ~ 10 km^2^ (Mills and Knowlton, 1991; Hibler, 1977) which is approximately the size of the largest islands in APIS. Home range size reported for red fox was smaller and in 1 *∼*10 km^2^ scale (Ables, 1969; Dekker et al., 2001; Trewhella et al., 1988) and was smaller in size than some individual islands in APIS. Together with mainland distance-dependency, spatial correlation patterns of coyote and fox were consistent with our knowledge of their movement ability, i.e., coyotes have stronger dispersal ability and larger home ranges, thus, weaker distance-dependency and stronger intra-island spatial autocorrelation compared with foxes. Patterns of coyotes and foxes demonstrate that species are not interchangeable (Island Biogeography Theory) and that distance-dependency is modified by life history characteristics.

The fisher-marten system also had a spatial auto-correlation effect. Furnas et al. (2017) reported a meta-analysis on home range sizes in California, USA. Their results showed that home ranges of female fishers varied and were approximately 6 km^2^ within 20 km of the coast while approximately 13 km^2^ at 120 km from the coast. Male fishers had home range sizes that varied from 12 km^2^ at 20 km to approximately 27 km^2^ at 120 km to the coast. We did not detect strong intra-island spatial autocorrelation in fishers, which may indicate a relatively small home range for these animals on islands compared to other studies, which was consistent with prior knowledge that fisher home ranges decline when close to coasts (Powell, 1982; Yaeger, 2005). Studies of marten home ranges in Canada indicated their home ranges can vary from 10 ∼100 km^2^ (Smith and Schaefer, 2002), we also detected spatial auto-correlation for martens.

Further home range and movement studies may be needed to confirm our findings based on their distribution in these islands. Coyote and fox showed different patterns compare with studies on the mainland. Studies in Canada on the interaction between coyotes and foxes showed that they typically partition through habitat use. But this pattern depended heavily on prey abundance (Theberge and Wedeles, 1989). Evidence also showed coyotes may aggressively kill red foxes (Gese et al., 1996). The positive correlation between foxes and coyotes may suggest that foxes trade-off predation risk for prey availability in a prey-limited system (Mallinger et al., 2021). Given the relatively small scale of this island system, the environment was rather homogeneous and the only environmental variation we were able to obtain was the distance to the mainland which is, itself, spatial. The model has the ability to incorporate additional environmental variation but measurements were unavailable in the case study (e.g. prey abundance, habitat structure, etc.). To better understand the system with the proposed model, detailed environmental variables like prey biomass should be measured to further explain the positive correlation pattern. Further study should be conducted to further evaluate these spatial distributions.

## 5: Conclusion

We implemented a MRF-based multi-specific occupancy model that can account for both spatial auto-correlations and inter-specific interactions simultaneously. Our model explicitly incorporates interaction terms between different species together with environmental predictors, and spatial auto-correlations. Our model parameters have clear interpretations as log odds of presence of one species at a site change when *holding all other species/sites’ occupancy constant* which is not clear in some other types of multi-species occupancy models. For instance, the centering term used in Hepler et al. (2018) made the coefficient of environment confounded by interactions while multinomial regression in Rota et al. (2016) had no explicit interaction term. Our model requires similar data as other types of occupancy models, i.e. repeat detection history and a pre-defined adjacency matrix of spatial-autocorrelation. Our simulation showed the model should work well in a study from 10 × 10 = 100 sites to 25 ×25 = 625 sites. For smaller grids, we may lose some ability to identify spatial auto-correlation. By formulation, the model can incorporate more species and a larger site number at the cost of computing time.

We demonstrate our technique in a case study on two pairs of presumably competing species of carnivores in the Apostle Island National Lakeshore (Wisconsin, USA). We detected a partitioning pattern of fisher and marten occupancy, which can be explained by a flipped source-sink pattern in the islands. However, additional evidence from movement data or genetics might be needed to further confirm this observation. We detected a positive association between coyote and fox, which, is different from studies on mainland systems and deserves further study.

## Acknowledgment

We appreciate helpful discussions with Dr. Claudia Solís-Lemus from Wisconsin Institute for Discovery during manuscript preparation and data collection support from the National Park Service. We thank the many students and staff that have participated in data collection. Y.S. acknowledges Jialong Jiang and Binxu Wang’s discussion during model and algorithm development. This research was funded by the United States National Park Service (USDI) through the Apostle Islands National Lakeshore (GLNF CESU Agreement P14AC01180), NASA Earth and Space Science Fellowship (Grant number NNX16AO61H). Additional funding came from the University of Wisconsin-Madison’s Beers-Bascom professorship and the UW-Madison Department of Forest and Wildlife Ecology’s A. W. Schorger fund to TVD and Sigurd Olson Professorship in the Department of Natural Resources at Northland College to E.O. We greatly appreciate the helpful feedback from three anonymous reviewers on the early draft of this manuscript.

## Supplementary Materials

### S1: Stepping Stone Graph

**Fig. S1.**
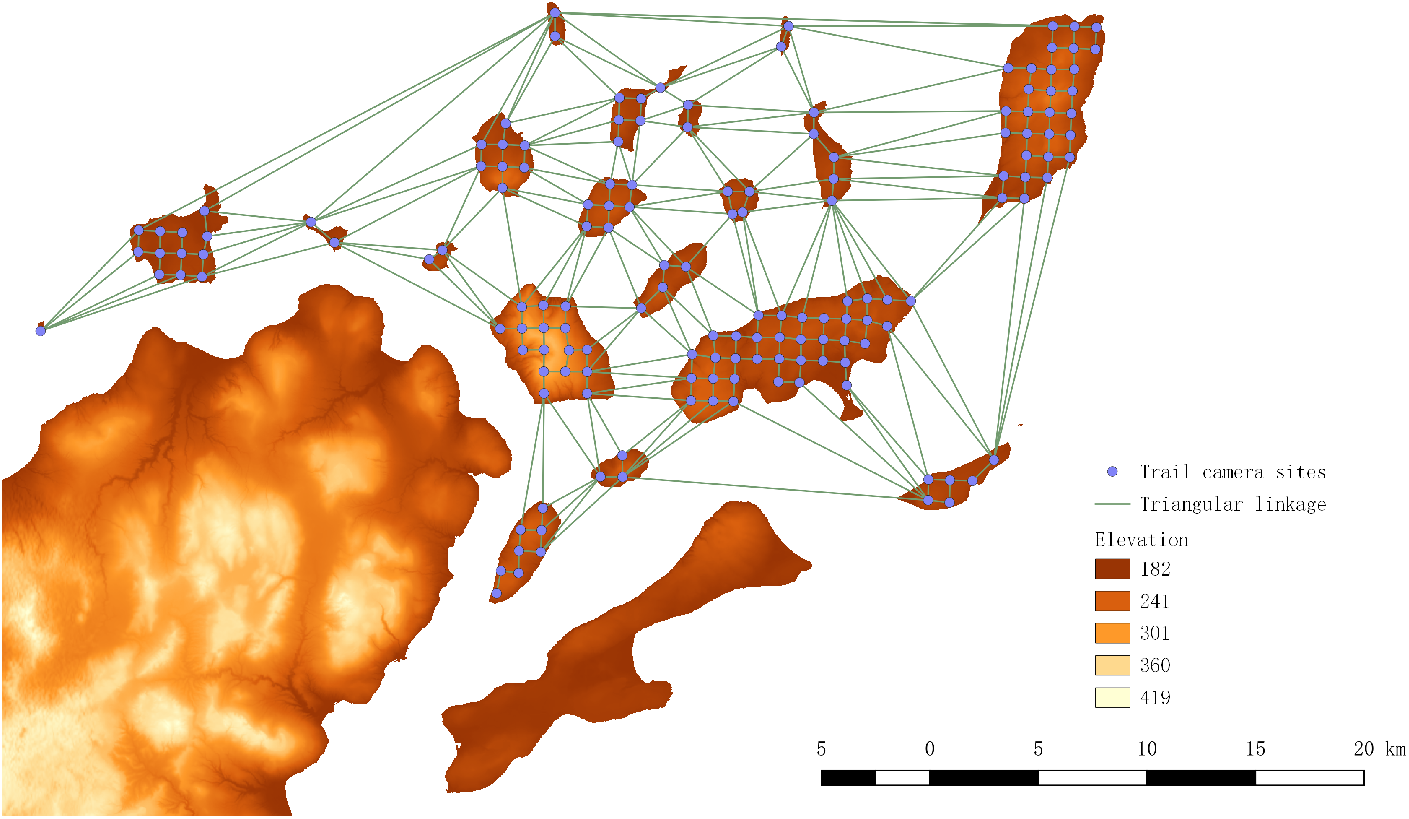
Graph used in Stepping stone model.

### S2: MH Algorithm and Bayes factor

Before Moller’s work in 2006, Bayesian inference on MRF models was precluded because the intractable normalizing constant is a function of parameters of interest. Moller proposed an auxiliary variable based on the Metropolis–Hastings ratio (Møller et al., 2006). A further development due to Murray et al. (2012) is called the single parameter change method. In our case, we follow Murray et al. (2012) by sampling an auxiliary variable *X* to follow the MRF distribution whose parameters are proposed *θ*^*Ⅎ*^. Together with imperfect detection, the Metropolis–Hastings ratio is given by eqn.S1.

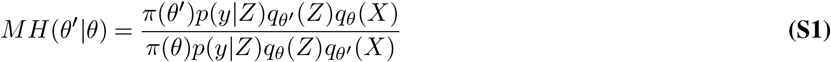

The sample can be drawn using the Coupling From the Past algorithm (CFTP Propp and Wilson (1996)) or long enough Gibbs chain for approximation. We tested the difference between using a perfect sample taken by CFTP algorithm and a Gibbs sample in the single parameter exchange algorithm. Results showed that if iteration for Gibbs is large enough (e.g. *>* 150) the posterior distribution sampled by these two methods were essentially the same (see example of Fig.S3). CFTP and Gibbs were implemented in `R` and `C++` modified from the `R` package `IsingSampler` (Epskamp, 2015) with help of `RcppArmadillo` (Eddelbuettel and Sanderson, 2014) and sparse matrix `C++` class provided by `R` package `Matrix` (Bates and Maechler, 2019) and `Armadillo` optimized for the sparse graph as we have (open sourced as `R` package `SparseIsingSampler` available on `GitHub`).

Posterior sample of **Z** will also be taken using a Gibbs algorithm, with fully conditional odds of being +1 as

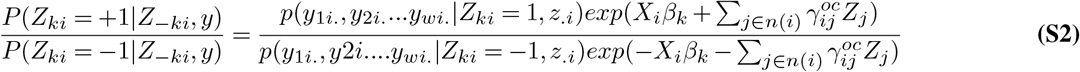

To calculate the Bayes factor, we need to calculate the likelihood of each sample using the harmonic rule (Raftery et al., 2006). To calculate the likelihood, we take a sample **Y** from a pre-specified parameter setting *ϕ*, the ratio of normalizing constant *C*(*θ*) and *C*(*ϕ*) can be calculated as the expectation: 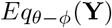. We can calculate log likelihood added by −*log*(*C*(*ϕ*)) which is intractable. However, by choosing the same *ϕ* for two competing models, we can calculate BF of two models by canceling out the intractable constant induced by *ϕ*.

### S3: Simulation

**Table S1.** Simulation setting for regular lattice.

| Parameter | Competition | No interaction | Sorting |
| --- | --- | --- | --- |
| $\beta_1$ | 0.5 | 0.5 | 0.3 |
| $\beta_2$ | 0.5 | -0.5 | 0.3 |
| $\eta^{in}$ | 0.25 | 0.25 | 0.25 |
| $\gamma_{oc}$ | -0.3 | 0 | 0.3 |
| $\gamma_{det}$ | -0.2 | 0 | 0.2 |

**Table S2.** Simulation setting for APIS.

| Parameter | Competition | No interaction | Sorting |
| --- | --- | --- | --- |
| $\eta_1^{ex}$ | 1 | 1 | 0.3 |
| $\eta_2^{ex}$ | 1 | -1 | 0.3 |
| $\eta^{in}$ | 0.2 | 0.2 | 0.2 |
| $\gamma_{oc}$ | -0.3 | 0 | 0.25 |
| $\gamma_{det}$ | -0.2 | 0 | 0.2 |

### S4: MCMC Diagnostic

We showed the trace plot and auto-correlation function of the interspecific interaction strength in occupancy *γ*_*oc*_ as an example diagnostic

**Fig. S2.**
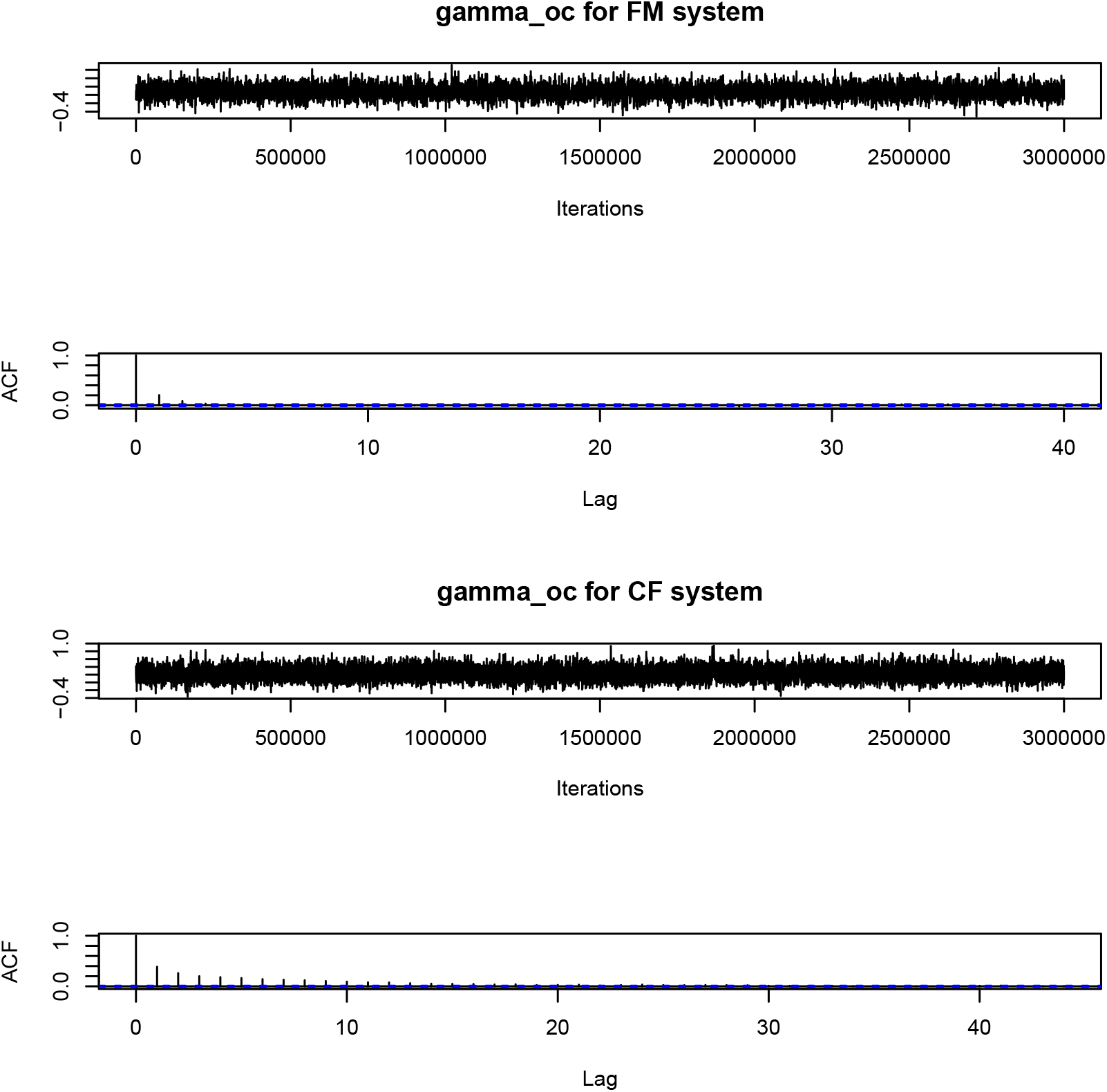
MCMC for *γ*_*oc*_ in FM and CF.

**Fig. S3.**
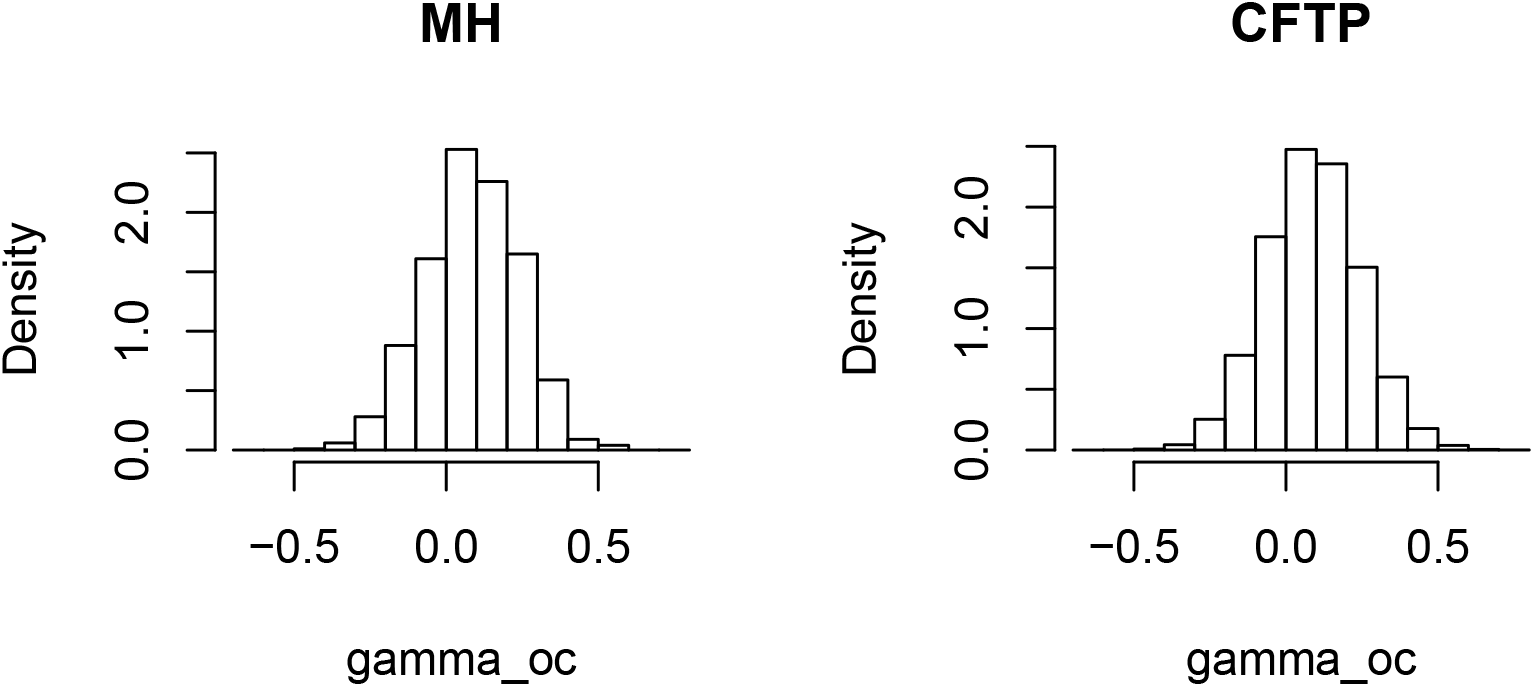
One example of same task using CFTP and MH, KS test had p-value of 0.9383.

### S5: Release of Prior on Detection Rate

Again it is not necessary to set this prior in a more general setting. But in the APIS case study, if we release this prior on the intercept of detection rate, the posterior had multiple modes for fisher and marten, due to the fact that fisher’s low naive detection can be due to both low occupancy or low detection, these results won’t influence our conclusion about the fisher and marten system but rather influence the numerical result of spatial auto correlation strength. This will cause a very large and unrealistic mainland-island strength in fisher S4.

**Fig. S4.**
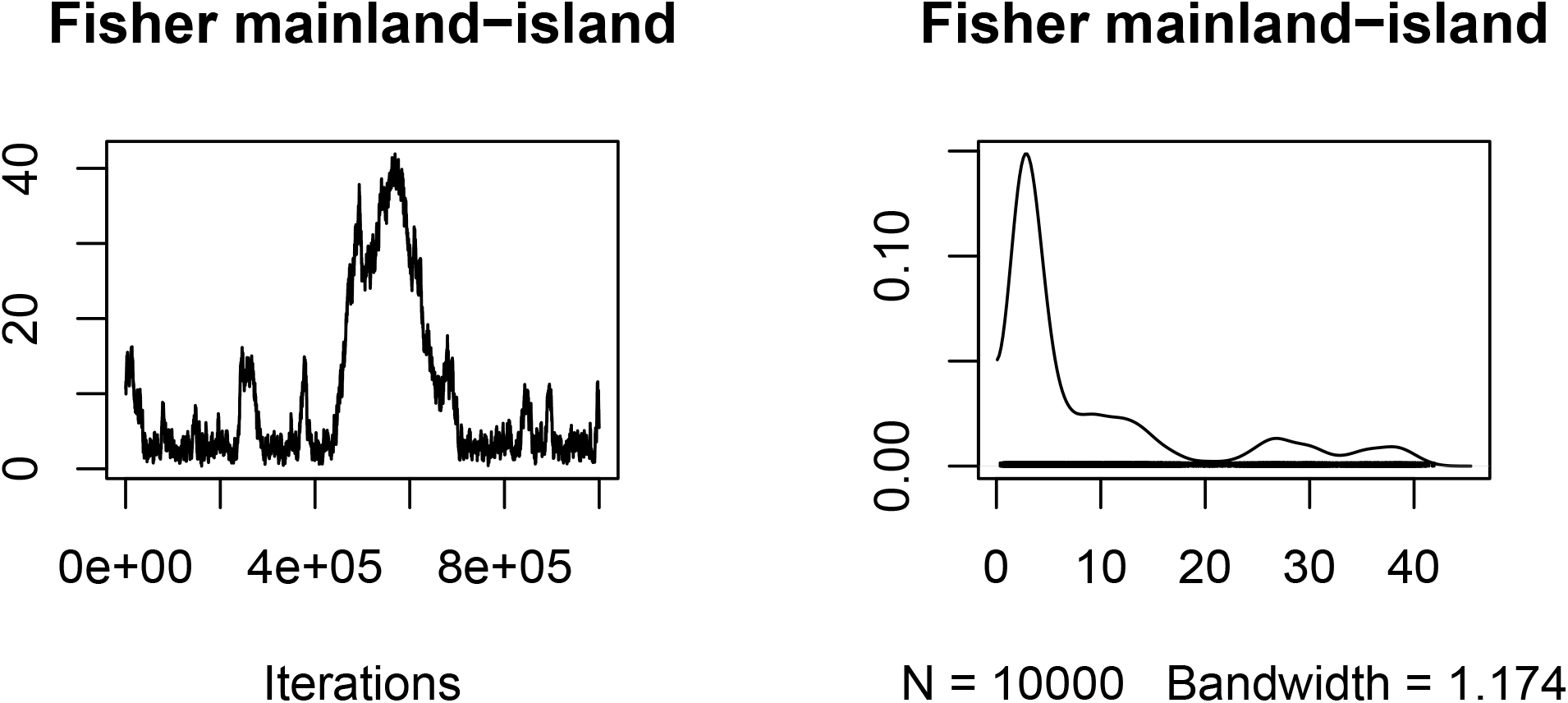
Multiple modes in fisher’s mainland-island strength if release the prior.

